# Respiratory suppression enhances voluntary control of human action

**DOI:** 10.64898/2026.01.15.699821

**Authors:** Haruna Miwa, Yusuke Akiyama, Jun Kunimatsu

## Abstract

Athletes often regulate their breathing to optimize motor performance, suggesting a close link between respiration and movement control. Previous studies have shown that respiration modulates neuronal oscillations and influences behavior. However, how respiratory control directly affects motor behavior remains unclear. Here, we demonstrate that breath holding during the preparatory period facilitates voluntary movement. We examined respiratory patterns while participants performed two types of eye movement tasks: pro-saccade and anti-saccade tasks. In the anti-saccade task, participants were required to suppress a reflexive saccade toward a visual target and instead generate a voluntary saccade in the opposite direction. To assess whether motivation modulates respiratory control, the tasks were conducted under two motivational conditions. Most participants suppressed their breath during the anticipatory period preceding saccade initiation, regardless of task type, and this tendency was more pronounced under high motivation. In the anti-saccade task, stronger respiratory adjustments were associated with shorter reaction times, whereas no such relationship was observed in the pro-saccade task. Moreover, intentional breath holding during the preparatory period selectively reduced reaction times in the anti-saccade task. These findings indicate that individuals spontaneously hold their breath to facilitate voluntary actions in highly motivated situations, and that this effect can be reproduced through intentional breath holding. We propose that anticipatory breath holding may play a functional role in proactively modulating neural processes underlying voluntary movement control.

## Introduction

Athletes regulate their breathing to optimize motor performance (Wells and Norris, 2009; HajGhanbari et al., 2013; Migliaccio et al., 2023). Although the beneficial effects of breathing control have been widely recognized from long-term experience, the underlying mechanisms remain poorly understood.

Breathing is a fundamental physiological function essential for sustaining life through gas exchange, and it is one of the few autonomic functions that can be voluntarily controlled. Because of these unique properties, breathing has also gained attention as a regulatory system that flexibly responds to internal and external conditions and interacts bidirectionally with cognitive processes (Takahashi et al., 2005; Russo et al., 2017). Previous studies have shown that respiration influences behavior by modulating neural oscillations (Tort et al., 2018; Folschweiller and Sauer, 2021). This synchronization is suggested to influence neural information processing and behavioral performance. Indeed, during inspiration, cognitive functions such as perception, attention, and memory are enhanced (Beh and Nix-James, 1974; Gallego et al., 1991; Perl et al., 2019; Molle et al., 2023), whereas during expiration, the frequency of motor actions such as button presses increases (Park et al., 2020). Furthermore, human participants tend to synchronize their breathing with predictable events and the timing of movements (Tobin et al., 1986). This tendency has been consistently observed across a wide range of cognitive tasks and shown to be reproducible across trials (Harting et al., 2025). These observations raise the possibility that respiratory control may constitute a component of proactive regulation of internal state.

It is known that the proactive process involves preparing attentional, perceptual, and action-related processes in advance to optimize performance in anticipation of upcoming events (Braver, 2012). Previous studies reported that the proactive process is particularly evident before motor actions that require voluntary control, including self-initiated movements (Maimon and Assad, 2006; Kunimatsu and Tanaka, 2012; Kunimatsu et al., 2019) and response inhibition (Meyer and Bucci, 2016; Liebrand et al., 2017). The anti-saccade task has been widely used to study the neural mechanisms of voluntary and inhibitory control of movement (Hallett and Adams, 1980; Rayner, 1998; Hutton, 2008; Riek et al., 2023). Ramping neuronal activity preceding anti-saccade onset has been observed in several cortical and subcortical regions, and this proactive signal has been shown to play a critical role in correct performance (Kunimatsu and Tanaka, 2010; Kunimatsu et al., 2016).

The neural circuits underlying saccadic eye movements have been characterized in considerable detail which makes the paradigm suited for interpreting how respiratory modulation influences motor behavior at the neural level. Furthermore, saccades impose minimal motor demand and do not involve body movement or postural changes. This allows for repeated execution without substantially altered breathing patterns. Thus, oculomotor tasks are particularly suitable for investigating respiratory control for movement. Additionally, saccadic eye movements are known to be facilitated by increased motivation (Takikawa et al., 2002; Watanabe and Hikosaka, 2005). Elevated motivation enhances proactive control and promotes internal state adjustments before movement (Qiao et al., 2018). Therefore, if breathing forms part of the proactive process, respiratory control is expected to be modulated by motivational state. In this study, we investigated spontaneous respiratory control during an anti-saccade task under different motivational states, and its effect on motor performance.

## Materials and Methods

### Participants

Forty-six healthy individuals participated in this study. Participants were randomly assigned to one of two experimental groups. In Experiment 1 (n = 30; 11 females, 19 males; mean age = 22.5 ± 2.5), participants initiate each trial when they pressed a button (SR-5 r1, LabHackers Research Equipment Inc.) over 2000 ms after previous trial, whereas in Experiment 2 (n = 16; 8 females, 8 males; mean age = 26.1 ± 8.1), each trial started automatically without a button press 4,000–6,000 ms after previous trial. All participants were naive to the purpose of the experiments and had no prior experience with similar tasks to prevent potential confounding effects from previous exposure. This study was approved by the Institutional Review Board of the Faculty of Medicine at the University of Tsukuba (1710-1). All procedures were conducted in accordance with the approved regulations. All participants provided written and verbal informed consent.

### Visual stimuli and behavioral tasks

We used two saccade paradigms: a pro-saccade task, in which participants made a reflexive saccade toward the visual target, and an anti-saccade task, in which they suppressed a saccade to the visual target (pro-saccade) and voluntarily made saccades away from it (Hallett, 1978). Participants were instructed to make a pro-saccade in response to the target when the fixation point was green, and an anti-saccade when it was red. Following a fixation period of 2,000–4,000 ms, a white target appeared 8° to the left or right of center on the screen, and participants made a saccade according to the task rule. In both saccade tasks, participants were required to move their eyes to the 3° “window” that surrounded the saccade goal within 2,000 ms of the target onset. In anti-saccade task, when participants successfully fixed within the correct window, a visual marker was presented to indicate that their gaze direction was correct, after which they were required to maintain their eye position for 300 ms. Upon successful saccade execution, a high-pitched tone (2000 Hz, 100 ms) was presented. When the saccade direction was incorrect, a low-pitched tone (200 Hz, 100 ms) was presented. If an error occurred during the trial (e.g., fixation break), an intermediate-pitched tone (400 Hz, 100 ms) was delivered via a speaker (MS-2, Marshall Amplification Ltd.).

In Experiment 1 (Fig. 1A), each trial was initiated with a white fixation point (0.6° square spot) when the participant pressed the button. Participants were required to maintain their eye position within 5° of the central fixation point for 600 ms. If a blink occurred during a period in which fixation was required, the fixation duration was extended by the blink duration. After an initial fixation period, the white fixation point changed color to either green or red. The saccade paradigms were conducted under two conditions to examine whether motivation affects respiratory control. In “feedback condition”, participants were instructed to perform the task as quickly as possible and received feedback (300 ms) based on their reaction time at the end of each trial. One of six words (“amazing”, “excellent”, “great”, “nice”, “good”, or “ok”) was presented depending on the participant’s reaction time (Fig. 1A, bottom). Feedback thresholds were determined based on the average reaction time (250 ms) with a fixed step width of 18.75 ms. The range from “amazing” (< 175 ms) to “good” (231.25–250 ms) was divided evenly at 18.75 ms intervals. The “ok” covered a wider range (250–287.5 ms), and no feedback was presented when the reaction time exceeded 287.5 ms. In “no-feedback condition”, participants were instructed to remain calm while performing the same task and received no such feedback.

**Figure 1.**
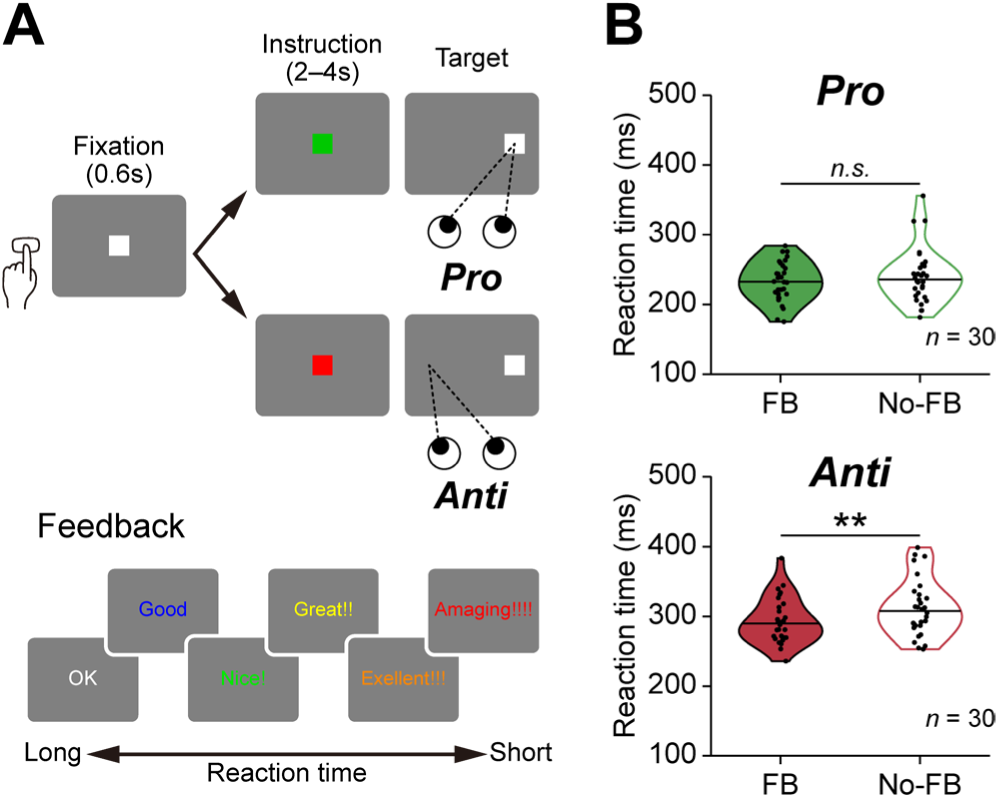
Two types of eye movement tasks. ***(A)*** Sequence of events in the pro-saccade and anti-saccade tasks. Following the button press that initiated each trial, a white fixation point appeared at the center of the screen. The task type was instructed by the color of the instruction cue (green, pro-saccade; red, anti-saccade). These tasks were conducted under two conditions: feedback (FB) and no-feedback (No-FB). In the feedback condition, one of six words was presented after the saccade depending on their reaction time. ***(B)*** Reaction times in the feedback and no-feedback conditions for the pro- and anti-saccade tasks. Violin plots show the distribution of participants’ mean reaction times, with individual means overlaid as dots. The horizontal lines indicate the mean values. (paired-samples Student’s t-test; \**p* < 0.05, \*\**p* < 0.01)

In Experiment 2 (Fig. 5A), each trial was started with a green or red fixation point after inter-trial interval (4,000–6,000 ms). Note that participants do not need to press the button. The verbal feedback was not used in this experiment.

### Experimental procedure

The experiment was conducted in a dark, sound-attenuated booth with participants seated in a chair. The participant’s head was stabilized using a head fixation device (HANDAYA Co., Ltd.). Participants wore a nasal cannula to measure respiration during the task. To ensure that they would breathe only through their noses, their mouths were sealed with surgical tape. Visual stimuli were presented on a 24-inch LCD monitor (ProLite E2483HS, iiyama Inc.; 59 Hz) positioned 47 cm in front of the eyes, subtending a visual angle of 60° × 36°. NIMH MonkeyLogic, a MATLAB-based system (MathWorks), was used for generating stimuli and controlling the behavior of the participants. At the beginning of the experiment, eye position calibration was performed by having participants fixate on a series of yellow squares presented sequentially on the screen. In both Experiments 1 and 2, the target direction (left or right) and saccade type (pro-saccade or anti-saccade) were pseudo-randomized within each session while ensuring an equal number of trials for each condition. Participants took a 1-min break after each session, and the break was extended if they reported fatigue or drowsiness. Each participant completed 10 practice trials before main trials.

In Experiment 1, each participant completed 10 practice trials and 40 main trials under each of the two conditions (feedback and no-feedback), including incorrect trials. A total of 80 main trials were collected for analysis.

In Experiment 2, each participant performed the task under two conditions to examine whether intentional breath holding affects task performance. In the “breath-hold” condition, participants were instructed to intentionally hold their breath during instruction period (2,000–4,000 ms). In the “control” condition, participants performed the task while breathing normally. Each participant completed 10 practice trials and continued the main task until they achieved 40 correct trials under each of the two conditions (breath-hold and control). This was done to collect reaction time data needed to analyze the relationship between respiration and task performance in Experiment 2. A total of 80 correct main trials were collected for analysis.

### Data acquisition and analysis

Respiration was measured as nasal airflow using a nasal cannula (PowerLab, ADInstruments). Eye movement and pupil diameter data were recorded using an eye-tracking system (iRecHS2, AIST). These data were sampled at 1 kHz and recorded during the experiments. All off-line analyses were performed using MATLAB (MathWorks).

Respiratory signals were band-stop filter between 1 and 500 Hz to reduce noise. After filtering, the signal was binarized using a predefined threshold, with values above the threshold classified as inspiration and values below as expiration. For each experimental condition, peaks of inspiratory and expiratory signals were detected, and the average peak values were calculated. Based on these averages, the signal was clipped from above for inspiration and from below for expiration, and each component was normalized separately. The normalized inspiratory and expiratory signals were then combined and further normalized to quantify the overall magnitude of respiratory fluctuations for each experimental condition.

Eye movement signals were low-pass filtered with a cutoff frequency of 60 Hz to remove high-frequency noise. Saccades were identified when horizontal eye displacement exceeded 5°. In trials in which a saccade was detected, saccade latency was defined as the time from target presentation to the earliest time point at which horizontal eye displacement exceeded 1.5°, preceding the detected saccade. In Experiment 2, only trials with reaction times ≤ 400 ms for the pro-saccade task and ≤ 500 ms for the anti-saccade task were included in the analysis.

Pupil diameter signals were preprocessed to remove blinks and artifacts as follows. First, abrupt changes in the pupil diameter signal were detected using the temporal derivative, and these samples were treated as outliers and marked as missing values. In addition, consecutive periods with extremely small derivative values were identified as signal loss associated with blinks, and samples in these periods, including the surrounding time points, were also marked as missing. The pupil diameter signal containing missing values was temporarily reconstructed using linear interpolation, after which a low-pass filter with a cutoff frequency of 5 Hz was applied to remove high-frequency noise. After filtering, samples corresponding to the original missing intervals were set back to missing values.

For each trial, two-time windows were defined; a baseline period (−1,500 to −500 ms from fixation onset in Experiment 1 and instruction onset in Experiment 2) and an anticipatory period (from 1,800 ms after instruction onset to target onset). To quantify respiratory fluctuations within these periods, the absolute difference between each sample of the normalized respiration signal (𝑥_𝑖_) and the midpoint of its range (0.5) was calculated and averaged across all samples within the defined period. This value was used as the respiratory change index, defined as:

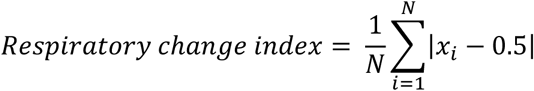

where N is the number of samples within the target period, and 𝑥_𝑖_ is the value of each sample in the normalized respiratory signal. In addition, pupillary fluctuations within the same time windows were quantified by averaging the preprocessed pupil diameter signal across all samples in each period, yielding a pupillary change index.

## Results

### Behavioral effects in task conditions

This study aimed to clarify how respiratory adjustments, particularly during the event anticipatory period, contribute to motor performance. We employed a saccade task consisting of pro-saccade and anti-saccade (Fig. 1A, top), allowing comparison of respiratory modulation between reflexive and voluntary movements requiring inhibitory control. In Experiment 1, participants performed saccade tasks under feedback (FB) and no-feedback (No-FB) conditions. The verbal feedback based on reaction time was provided at the end of each trial (Fig. 1A, bottom). Previous studies have shown that receiving performance-related feedback increases subjective motivation and is accompanied by shorter reaction times (Eckner et al., 2011). Based on these reports, the FB condition was expected to induce a higher motivational state compared with the No-FB condition. Actually, reaction times in the anti-saccade task were significantly shorter in the FB condition than in the No-FB condition (Fig. 1B; paired-samples Student’s t-test, *p* = 2.8 × 10^-3^), but they were not significant in the pro-saccade task (*p* = 0.088). These results suggested that participant performed the tasks, at least anti-saccade task, with high motivation in the FB condition relative to the No-FB condition.

### Patterns of breathing during the pro- and anti-saccade task

To examine how respiration changed in relation to task events within each trial, we first analyzed the temporal dynamics of respiratory phase during task execution. Figure 2A shows data from a representative participant, in which trials are categorized by task types (pro- and anti-saccade) and conditions (FB and No-FB), and respiratory phase is visualized as a heat map. Each row represents a single trial, aligned to the onset of the instruction, and trials are sorted by the duration of the randomized fixation period between instruction and target onset. Warmer colors indicate inspiration, cooler colors indicate expiration, and values near green represent minimal respiratory change, corresponding to breath holding. This data revealed that participants suppressed their breathing during the instruction period, while anticipating target onset.

**Figure 2.**
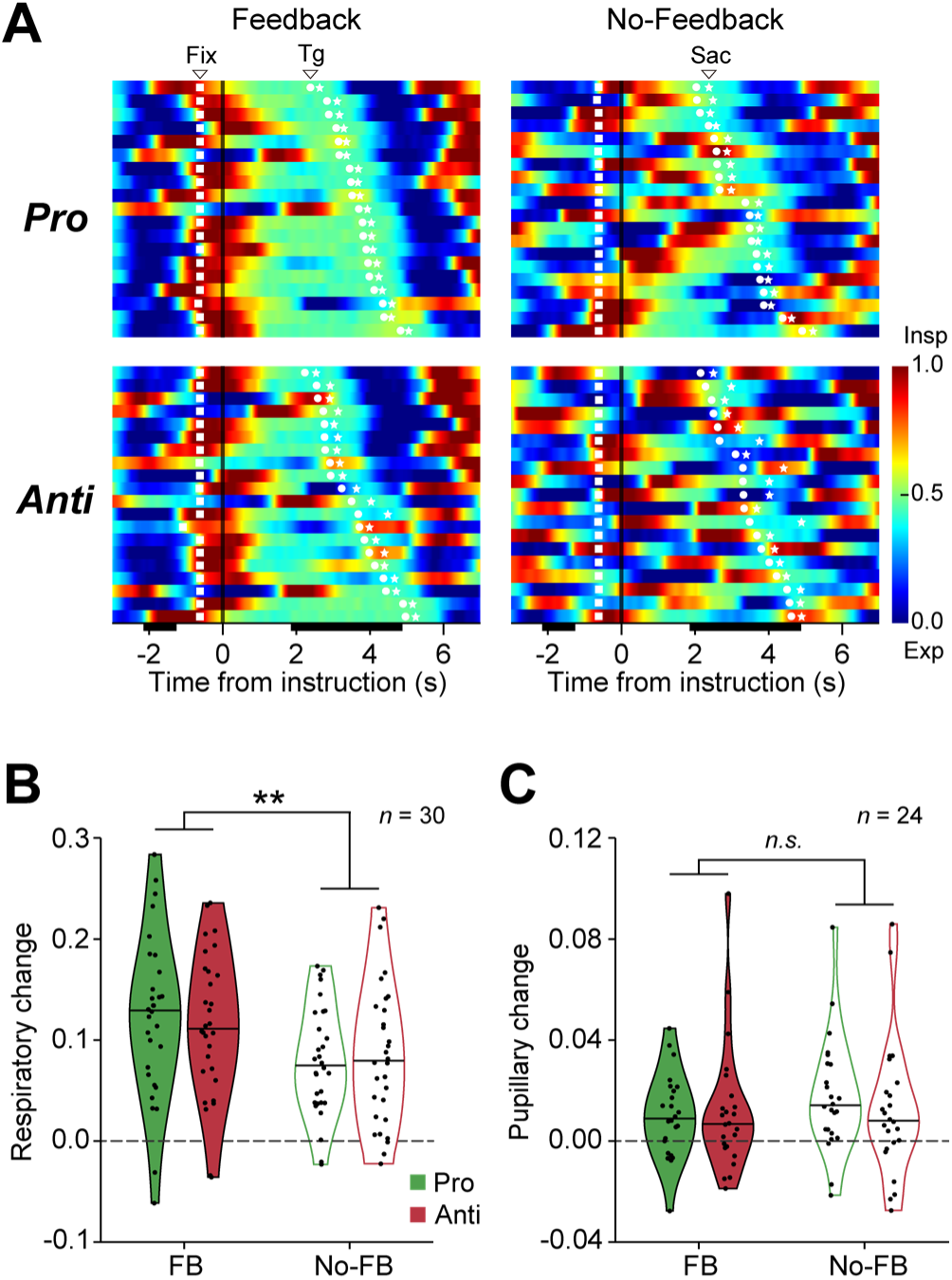
Task-related respiratory changes during anticipatory period. ***(A)*** Heat map of the respiratory waveform from a representative participant. Each row represents one trial, with red showing inhalation and blue showing exhalation. The squares show the fixation cue, the circles show the target, and the stars show the saccade onset. Trials are aligned to the timing of the instruction (vertical black bar) and sorted by the duration between the instruction and the target. ***(B, C)*** Respiratory change ***(B)*** and pupillary change ***(C)*** in each condition for the pro- and anti-saccade tasks. Violin plots show the distribution of participants’ mean changes, with individual means overlaid as dots. The horizontal lines indicate the mean values. (two-way ANOVA; \*\**p* < 0.01)

To quantitation analysis, we defined a “respiratory change index” for each trial to characterize respiratory modulation relative to baseline (see Methods). This index was computed as the difference in respiratory variability between the baseline period and the anticipatory period, thereby reflecting the extent of respiratory suppression during anticipation. Figure 2B indicates the mean values of the respiratory change index, computed separately for each task type and condition. Consistently across all task types and conditions, participants exhibited significant respiratory suppression relative to baseline (one-sample Student’s t-test, pro-saccade w/ FB, *p* = 1.8 × 10^-9^; anti-saccade w/ FB, *p* = 5.2 × 10^-10^; pro-saccade w/ No-FB, *p* = 3.9 × 10^-9^; anti-saccade w/ No-FB, *p* = 1.2 × 10^-7^). A two-way ANOVA (task: pro-saccade vs. anti-saccade; condition: FB vs. No-FB) further revealed a significant main effect of condition (*p* = 5.5 × 10^-3^), indicating that respiratory suppression was more pronounced in the FB condition, which was associated with a higher motivational state.

In contrast to the results of respiratory changes, the ANOVA did not reveal any significant effects in pupil diameter (*p* > 0.38). These results indicate that respiratory modulation during instruction period was not accompanied by corresponding changes in pupil diameter, an index of autonomic responses. This suggests that respiratory adjustment during the task arises from mechanisms distinct from autonomic system.

### Breathing in relation to task performance

We showed systematic respiratory suppression during the anticipatory period. If this modulation serves a functional role in behavior, it should be reflected in task performance. Accordingly, we analyzed the relationship between task-related respiratory changes and reaction time. Figure 3 (left) shows the correlation between the respiratory change index and reaction time for a representative participant (the same as in Fig. 2A), separately for each task type (pro- and anti-saccade). No clear correlation was observed in the pro-saccade task (*r* = -0.20, *p* = 0.21). In contrast, in the anti-saccade task, a significant negative correlation was found, indicating that trials with larger index values, reflecting stronger respiratory suppression, were associated with shorter reaction times (*r* = -0.34, *p* = 0.040). To assess whether this relationship was consistent at the group level, we compared the distributions of correlation coefficients across all participants (Fig. 3, right panel). In the pro-saccade task, the distribution of correlation coefficients showed no significant bias (Wilcoxon signed-rank test, *p* = 0.11), whereas in the anti-saccade task, the distribution was significantly shifted toward negative values (*p* = 0.023). These results reveal respiratory modulation during anticipatory periods is associated with shorter reaction times in the anti-saccade task.

**Figure 3.**
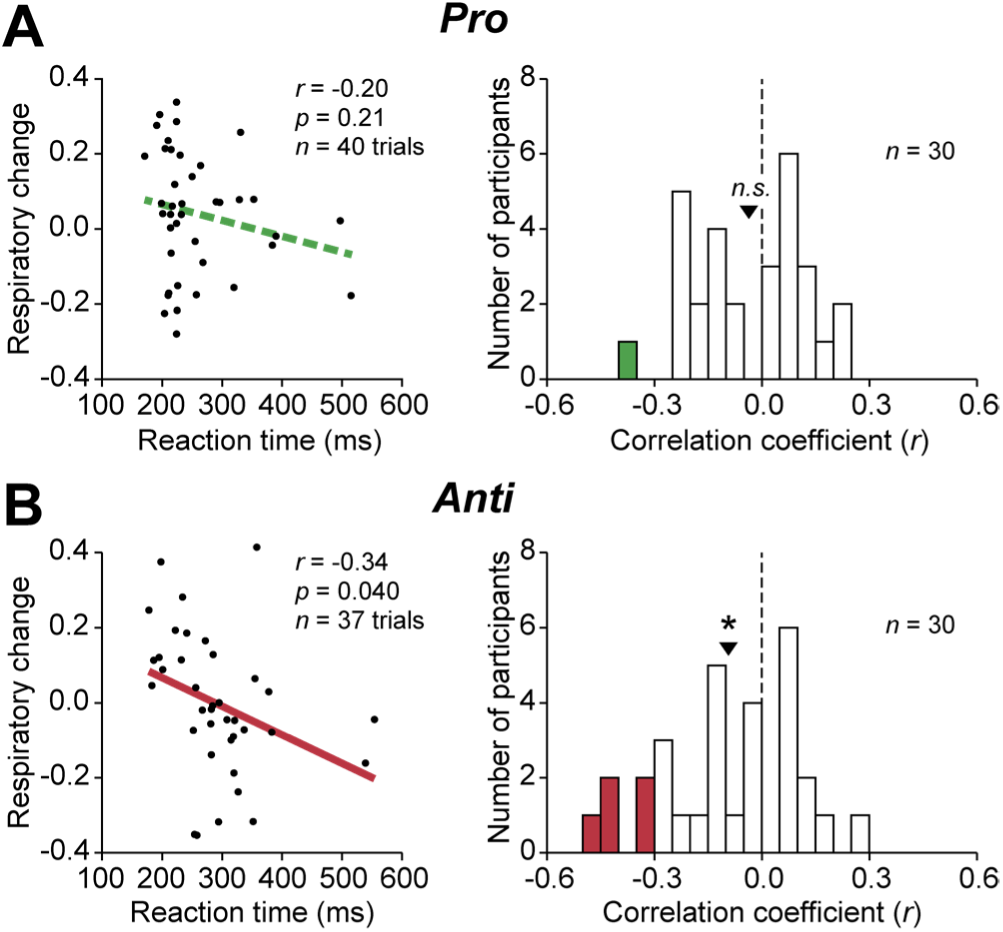
Relationship between respiratory change and task performance. ***(A, B)*** Scatter plots show the correlation for a representative participant (left), and histograms show the distribution of correlation coefficients across all participants (right) in the pro-saccade task ***(A)*** and in the anti-saccade task ***(B)***. In the scatter plots, green dashed lines indicate non-significant linear regression fits, whereas red solid lines indicate significant fits. In the histogram, participants showing significant correlations are marked in green or red, respectively (individual linear regression, *p* < 0.05). Triangles indicate the median of each distribution. (Wilcoxon signed-rank test; \**p* < 0.05)

Because previous studies reporting that respiratory phase (inspiration and expiration) differentially affects task performance including reaction time (Beh and Nix-James, 1974; Gallego et al., 1991; Tort et al., 2025), we examined whether the respiratory phase at target onset influenced reaction time. Figure 4 (left panel) summarized the respiratory phase across all participants using polar coordinates. Regardless of task type, the majority of target presentations occurred during the expiratory phase. In addition, we examined the relationship between respiratory phase at target onset and the respiratory change index. Trials in which the target was presented during expiration showed greater respiratory suppression than those presented during inspiration (Fig. 4, middle panel; paired-samples Student’s t-test, pro-saccade, *p* = 1.8 × 10^-4^; anti-saccade, *p* = 1.1 × 10^-3^). Next, we analyzed reaction time differences between trials in which the target was presented during inspiration and those during expiration. Change of reaction time was not observed between the two phases (Fig. 4, right panel; paired-samples Student’s t-test, pro-saccade, *p* = 0.76; anti-saccade, *p* = 0.16), indicating that respiratory phase at target onset did not influence reaction time in this experiment. These results indicate that although participants have adjusted the timing of their breathing to avoid entering the inspiratory phase at target presentation, respiratory phase was not a determining factor for reaction time. Moreover, this finding is consistent with the preceding our results showing that the magnitude of respiratory suppression affects performance, suggesting that breath holding has a greater impact on reaction time than respiratory phase.

**Figure 4.**
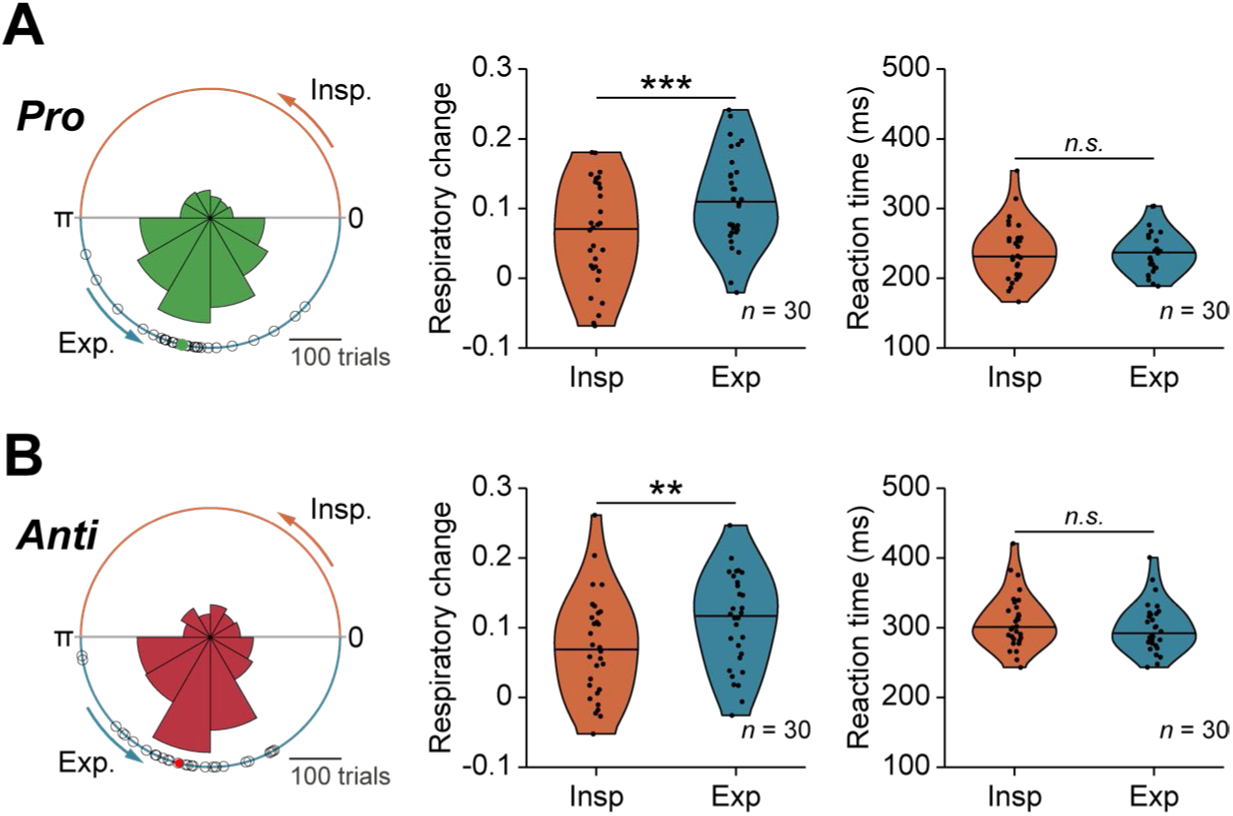
Effect of respiratory phase at target onset. ***(A, B)*** Polar histogram (left) shows the distribution of all the target onset from all participants in the pro-saccade task ***(A)*** and in the anti-saccade task ***(B)***. Empty black circles represent each participant’s mean respiration phase at target onset. The green or red dot indicates the grand-averaged respiration phase at target onset. Violin plots (middle) show respiratory changes, computed separately for trials in which the target appeared during inspiration or expiration, with individual participants’ means overlaid as dots. The horizontal lines indicate the mean values. (paired-samples Student’s t-test; \*\**p* < 0.01, \*\*\**p* < 0.001) Violin plots (right) show reaction times, computed separately for trials in which the target appeared during inspiration or expiration, with individual participants’ means overlaid as dots. The horizontal lines indicate the mean values. (paired-samples Student’s t-test; \**p* < 0.05)

**Figure 5.**
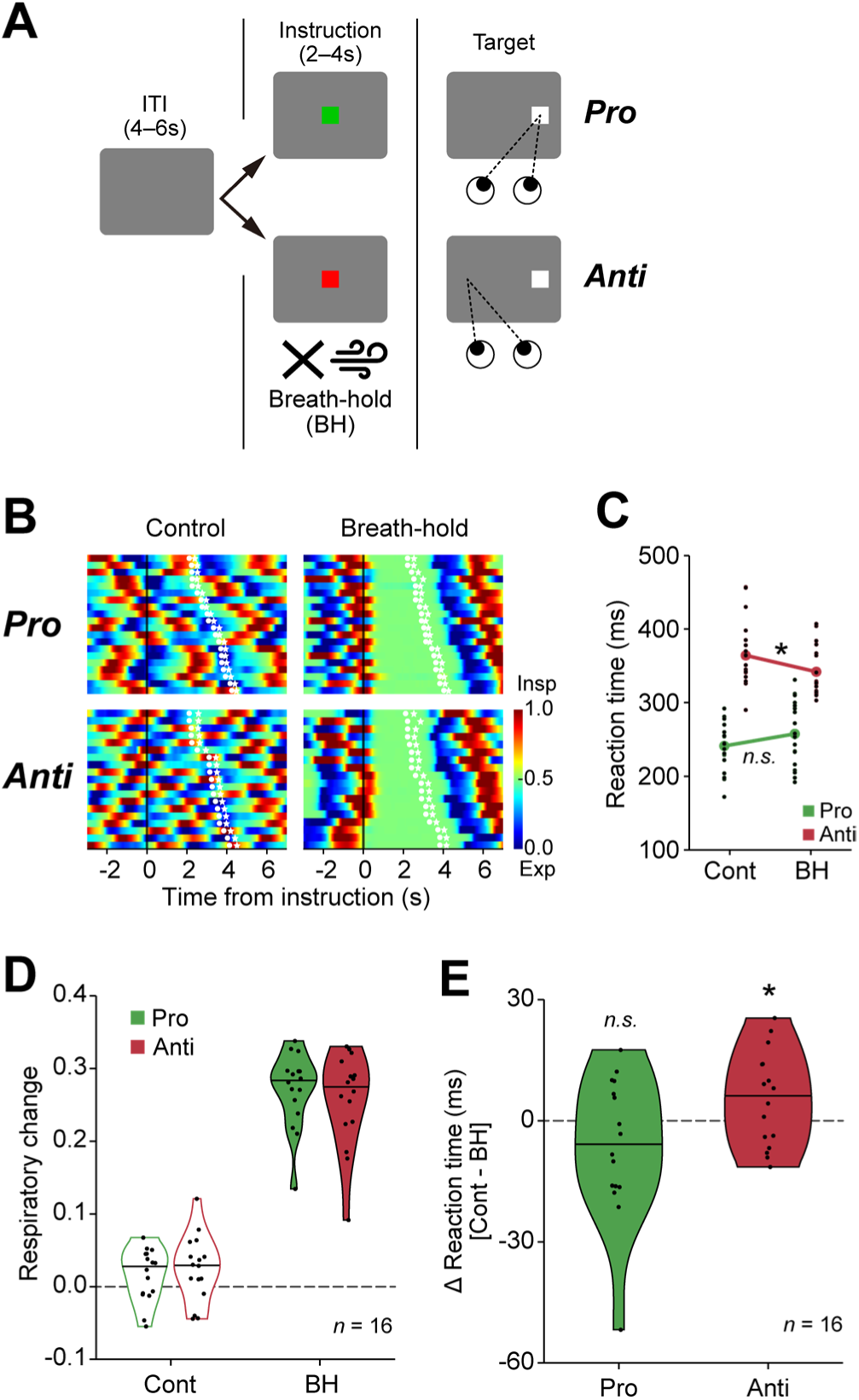
Performance under the intentional breath holding. ***(A)*** Sequence of events in breath-hold condition. After a 4–6 s inter-trial interval (ITI), each trial begins with the presentation of an instruction cue (green, pro-saccade; red, anti-saccade). Participants were instructed to fixate it and to intentionally hold their breath while the point was presented. ***(B***) Heat map of the respiratory waveform from a representative participant among conditions. ***(C)*** Reaction times for a representative participant in the pro- (green) and anti-saccade (red) tasks for each condition. Black dots represent reaction times on individual trials, and the colored dot represents the mean value (paired-samples Student’s t-test; \**p* < 0.05). ***(D)*** Respiratory change in each condition for the pro- and anti-saccade tasks. Violin plots show the distribution of participants’ mean changes, with individual means overlaid as dots. ***(E)*** Reaction time differences between the control and breath-hold conditions for pro- and anti-saccade tasks. Violin plots show the distribution of participants’ mean reaction times, with individual means overlaid as dots. The horizontal lines indicate the mean values. (one-sample Student’s t-test; \**p* < 0.05)

### Effects of intentional modulation of breathing

Although Experiment 1 demonstrated that spontaneous respiratory suppression observed during event anticipatory period was associated with improved task performance, the causal relationship remains unclear. Therefore, the task was modified to examine whether a similar effect would be observed when participants intentionally suppressed their breathing.

In Experiment 2, we used an intentional breath-holding condition, in which participants were instructed to maintain breath holding throughout the instruction period. This period corresponds to the interval in experiment 1 during which participants suppressed their breathing (Fig. 5A). Unlike Experiment 1, the button press was used to initiate each trial and the initial white fixation point was omitted. This manipulation was designed to prevent participants from predicting trial onset and to avoid inducing unintentional respiratory adjustments. Actually, the breath holding was not observed in the control condition, in which participants were instructed to breathe naturally (one-sample Student’s t-test, pro-saccade, *p* = 0.016; anti-saccade, *p* = 0.074), whereas respiratory suppression was confirmed only in the breath-hold condition (Fig. 5B; pro-saccade, *p* = 8.7 × 10^-15^; anti-saccade, *p* = 2.8 × 10^-12^). To assess the effect of intentional breath holding for behavior, reaction times were examined across conditions. The breath-hold condition selectively reduced reaction times in the anti-saccade task (Fig. 5C; paired-samples Student’s t-test, *p* = 0.016), with no significant effect in the pro-saccade task (*p* = 0.86). The same results were consistently observed at the group level across 16 participants (Fig. 5D, E; one-sample Student’s t-test, pro-saccade, *p* = 0.92; anti-saccade, *p* = 0.047). These results demonstrate that anti-saccade reaction time is facilitated not only through automatic and spontaneous respiratory suppression during the task, but also through intentional breath holding. These findings suggest that breath-holding is a key factor to improve behavioral performance in the anti-saccade task.

## Discussion

In this study, we revealed that respiration is suppressed during the motor preparation in alignment with trial events, particularly under highly motivated condition (Fig. 2). Furthermore, this temporary breath holding was associated with shorter reaction times in the anti-saccade task (Fig. 3), indicating its contribution to voluntary motor performance. In addition, this contribution was reproduced through intentional breath holding (Fig. 5), indicating that respiratory control serves as a factor enhancing motor performance. These results suggest that, irrespective of motivational state, respiratory suppression may enhance motor performance, with potential applications in improving sports performance and motor rehabilitation.

### Respiratory modulation on behavioral control

The present study demonstrates that respiratory suppression during the anticipatory period is associated with improved performance in the anti-saccade task but not in the pro-saccade task (Figs. 2, 3). This effect cannot be readily explained by a general increase in arousal or alertness, as such non-specific factors would be expected to facilitate both pro- and anti-saccade performance. Instead, the selective effect observed in the anti-saccade task suggests that respiratory suppression is specifically related to processes required for inhibitory control and voluntary action selection. Consistent with this interpretation, Experiment 2 showed that intentional respiratory suppression was sufficient to reproduce the behavioral enhancement observed in Experiment 1, providing converging evidence for a functional contribution of respiratory suppression to task performance.

Previous studies have reported that respiratory phase synchronizes with neural oscillations, and it modulates motor and sensory information processing (Tort et al., 2018; Folschweiller and Sauer, 2021). In particular, in tasks in which stimulus timing is predictable, participants synchronize their breathing with trial events, and respiratory changes preceding responses have been shown to contribute to enhanced performance (Kluger et al., 2021; Harting et al., 2025). These findings support the interpretation that the respiratory control observed during the motor preparation period in this study contributes to performance enhancement by the regulation of internal state prior to motor execution.

Most previous studies have primarily focused on respiratory phase and have revealed that different phases have distinct effects on task performance (Beh and Nix-James, 1974; Gallego et al., 1991; Perl et al., 2019; Park et al., 2020; Molle et al., 2023). However, it has been suggested that the relationship between respiratory phase and reaction time depends on the state of respiratory control. Indeed, phase-dependent effects on reaction time have been observed when participants voluntarily control their breathing, whereas no such effects have been reported under natural breathing (Li et al., 2012). In this study, no instructions about breathing during task performance were provided. Therefore, the result of no clear phase-dependent effect on reaction time (Fig. 4) is consistent with these previous findings.

Because breathing control is modulated by the autonomic nervous system, it was plausible that the performance enhancement due to respiratory suppression observed in this study may involve modulation of autonomic activity. Pupil diameter is widely used as an index of autonomic nervous system activity (Grujic et al., 2024), and therefore we also analyzed pupil responses. However, no differences in pupil diameter were observed between conditions, and no significant correlation was found between pupil diameter and reaction time (Figs. 2C, 4). These results indicate that the performance enhancement associated with respiratory suppression cannot be explained by autonomic changes reflected in pupil diameter. In addition, pupil diameter is known to reflect changes in arousal and attention (van der Wel and van Steenbergen, 2018; da Silva Castanheira et al., 2021), suggesting that the reaction time modulation observed in this study was not mediated by these factors.

One possible mechanism by which breathing influences other motor behaviors is through changes in posture and body movements (Caron et al., 2004), and breath holding may suppress body movements and thereby enhance the performance of other motor actions. However, this study showed that the effect of breathing is not on limb movements, but on eye movements, which are thought to have less benefit in stopping body movement. In addition, participants’ head was stabilized during the experiment, minimizing the influence of body movement. Despite this, a significant relationship between respiratory suppression and reaction time was observed, suggesting that respiratory suppression influenced neural circuits involved in the control of voluntary movements.

### Mechanisms underlying the enhancement of anti-saccade performance

What mechanisms underlie the respiratory suppression and the enhancement of voluntary movement observed? Respiratory suppression during the motor preparation period was more pronounced in the FB condition with higher motivation (Fig. 2B). Previous studies repeatedly demonstrated a facilitative effect of motivation on motor performance (Wulf and Lewthwaite, 2016). In particular, motivational manipulations such as reward and task difficulty primarily affect the motor preparation state (Mir et al., 2011; Wilhelm et al., 2021). The basal ganglia are thought to play an essential role in adjusting motor preparation based on motivational context. These are considered to modulate motor gain, including movement speed and amplitude, according to motivational factors such as reward and task demands (Desmurget and Turner, 2010; Turner and Desmurget, 2010; Panigrahi et al., 2015). Similar modulation of motor preparation by motivational factors has also been reported in oculomotor tasks (Hikosaka et al., 2006; Larry et al., 2024). Based on these findings, it is possible that the basal ganglia were also involved in the motivation-dependent respiratory suppression observed during the motor preparation period. Consistent with this interpretation, stimulation of the caudate nucleus has been shown to induce apnea, indicating that the basal ganglia can influence respiratory activity in addition to motor control (Angyán, 1994). Taken together, these findings suggest that increased motivation may activate basal ganglia circuits, which in turn give rise to respiratory suppression as part of the adjustment of internal state during motor preparation.

Furthermore, in this study, respiratory suppression during the motor preparation period was not associated with reaction time in the pro-saccade task, but was associated with shorter reaction times only in the anti-saccade task (Figs. 3, 5). The anti-saccade task requires suppression of the automatic pro-saccade response toward the stimulus, followed by the generation of a voluntary eye movement in the opposite direction. Primate studies have shown that such control processes are known to involve the cortico–basal ganglia circuit comprising frontal and parietal cortices and the superior colliculus (Munoz and Everling, 2004; Kunimatsu and Tanaka, 2010; Jantz et al., 2017). In particular, neural activity in regions such as the dorsolateral prefrontal cortex (Johnston and Everling, 2006), frontal eye field (FEF) (Everling and Munoz, 2000), supplementary eye field (SEF) (Schlag-Rey et al., 1997; Amador et al., 2004), caudate nucleus (Watanabe and Munoz, 2011), and external segment of the globus pallidus (Yoshida and Tanaka, 2009) is more strongly enhanced during the motor preparation period prior to target presentation for anti-saccades than for pro-saccades (Everling et al., 1998; Everling and Munoz, 2000; DeSouza et al., 2003; Munoz and Everling, 2004). Pharmacological inactivation of some of these regions selectively increases error rates and delays reaction times in the anti-saccade task (Brown et al., 2007; Yoshida and Tanaka, 2009; Kunimatsu and Tanaka, 2010). These findings suggest that the respiratory suppression observed during the motor preparation period in this study contributed to the preparatory activation of inhibitory control networks associated with the anti-saccade task. Together, these findings suggest that respiratory suppression during action preparation reflects a global preparatory state that enhances the engagement of cortico–basal ganglia inhibitory control networks, thereby selectively facilitating voluntary action.

It has also been suggested that the neural substrates underlying respiratory control and voluntary motor control partially overlap. The dorsomedial frontal cortex, including the SMA, SEF, and pre-SMA, is known to be involved in voluntary control across multiple modalities (Tanji, 1996; Nachev et al., 2008; Kunimatsu and Tanaka, 2012). The SEF plays a key role in the planning and preparation of voluntary eye movements and supports the generation of anti-saccades through coordinated interactions with the FEF and the superior colliculus (Schlag-Rey et al., 1997; Amiez and Petrides, 2009; Stuphorn et al., 2010). The SMA has also been reported to be part of a cortico–subcortical network that is activated during voluntary breath holding (McKay et al., 2003), and afferent feedback from the respiratory system has been shown to modulate motor output, potentially by providing sustained excitatory input to the SMA (Laviolette et al., 2013). These findings suggest that respiratory control may influence motor preparatory processes via dorsomedial frontal structures.

Taken together, these findings suggest that increased motivation activates the cortico–basal ganglia circuit, including the SMA, leading to respiratory suppression during event anticipation as well as facilitation of voluntary movement. Furthermore, the finding from Experiment 2 that intentional breath holding enhanced motor performance raises the hypothesis that respiratory suppression may strengthen the activity of this circuit and thereby improve voluntary movement. From this perspective, respiratory suppression should not be regarded merely as a physiological byproduct but rather as a state variable that reflects or modulates the engagement of neural circuits underlying voluntary control. Future studies combining respiratory manipulation with neural recordings will be necessary to directly test the proposed neural mechanisms and determine whether respiratory suppression causally modulates activity in these circuits.

## Acknowledgments

We are grateful to E. Doi for valuable comments on this study. We also thank A. Yokoyama for their administrative support. This work was supported by grants from JST, PRESTO (grant numbers: JPMJPR21S4) and MEXT (21H05800 and 21H05036), and the Takeda Science Foundation.

